# The RNA demethylases ALKBH5 and FTO regulate translation of the ATF4 mRNA in sorafenib-treated hepatocarcinoma cells

**DOI:** 10.1101/2024.02.09.577647

**Authors:** Pauline Adjibade, Rachid Mazroui

## Abstract

Translation is one of the main gene expression steps targeted by cellular stress, commonly refereed as translational stress, which includes treatment with anticancer drugs. While translational stress blocks translation initiation of bulk mRNAs, it allows translation of a specific set of mRNAs known as short upstream open reading frames (uORFs)-mRNAs. Among these, ATF4 mRNA encodes a transcription factor that reprograms gene expression during various cellular stress towards functions required for cell response to stress. Stress-induced ATF4 mRNA translation occurs via a specialised mode that relies on the presence of uORFs upstream to the main ATF4 ORF. However, mechanisms regulating ATF4 mRNA translation, particularly towards chemoresistance, remained limited. Here, we report a role of both ALKBH5 and FTO, the two RNA demethylating enzymes in promoting translation of ATF4 mRNA in liver cancer Hep3B cells treated with sorafenib, a stress inducer used in chemotherapy. Depletion experiments confirmed that both enzymes are required for inducing ATF4 mRNA translation, while polyribosome assays coupled to RT-qPCR indicated that this induction of ATF4 mRNA translation occurs at its initiation step. Using *in vitro* methylation assays, we found that ALKBH5 is required for the inhibition of the methylation of a reporter ATF4 mRNA at a conserved adenosine (A235) site located at its uORF2, suggesting that ALKBH5-mediated translation of ATF4 mRNA involves demethylation of its A235. Preventing methylation of A235 by introducing an A/G mutation into the ATF4 mRNA reporter renders that reporter insensitive to ALKBH5 depletion, supporting the role of ALKBH5 demethylation activity in translation. Finally, targeting either ALKBH5 or FTO sensitizes Hep3B to sorafenib-induced cell death, contributing to their resistance. We concluded that ALKBH5 and FTO are novel factors that promote resistance to sorafenib treatment, in part by mediating translation of ATF4 mRNA.

## Introduction

Gene expression reprogramming is fundamental for cell growth, cell proliferation, and cell differentiation. Reprogramming of gene expression is also critical for cell adaptation to stress and survival, involving mRNA translational regulatory pathways^1,2^. Among these, phosphorylation of the translation initiation factor eIF2α is the main pathway that is induced during stress including endoplasmic reticulum (ER) stress, oxidative stress, nutrient deprivation, hypoxia, and cancer drug treatment^3,4^. The phosphorylation of eIF2α (PeIF2α) causes a global inhibition of protein synthesis concomitant with the formation of RNA stress granules (SG)^5^, composed of RNA and associated proteins (mRNPs)^5^. While these stress-induced SG are thought to represent a pool of mRNPs stalled in the process of translation initiation, contributing to PeIF2α-mediated global translation inhibition, PeIF2α itself promotes the preferential translation of specific mRNAs encoding stress.

ATF4 (Activating transcription factor) is a critical stress-induced factor whose expression relies on PeIF2α. ATF4 mRNA has been widely used as a PeIF2α-target paradigm, establishing that PeIF2α-mediated translation occurs via a specialized mode of translation initiation relying on the presence of short upstream open reading frames (uORFs) at 5’untranslated region (UTR) of mRNAs^4^. In absence of PeIF2α (i.e., no stress), high translation at uORFs prevents ribosomes from translating the overlapping ATF4 ORF^6,7^. Upon stress, PeIF2α allows ribosomes to bypass the presence of the inhibitory uORFs, thus initiating translation at the main ATF4 ORF, which then activates transcription of downstream genes encoding either apoptotic (e.g., ATF3 and CHOP), or adaptive (e.g., chaperones, antioxidants and amino acid synthases) factors^8,9^. These opposite functions of ATF4 in the cellular stress response depend largely on its expression level linked to its translational regulation by PeIF2α. In cancer, the level of ATF4 mRNA translation induced by PeIF2α is also critical to promote either cell death^8^ or survival^9^ to ER stress generated upon treatment with genotoxic drugs, requiring regulatory mechanisms. We have previously described one such mechanism involving the activity of the RNA helicase DDX3^10^. In this study, we showed that DDX3 promotes PeIF2α-induced ATF4 translation at the earliest step of its initiation by recruiting the translation initiation eIF4F factor to the cap structure of the 5’end of ATF4 mRNA. More recently, new mechanisms that promote PeIF2α-induced ATF4 mRNA translation were reported involving the activity of the RNA demethylases ALKBH5 and FTO in amino acid-starved mouse embryonic fibroblasts^11^, and possibly in human HEK293T cells treated with the ER stressor thapsigargin^12^.

The N6-methyladenosine (m^6^A) is a key chemical modification in RNA regulating multiple RNA-based processes and was recently reported as critical for stress- and cancer-regulated RNA metabolism including translation^13^. This m^6^A modification is usually present in the consensus [G/A m^6^AC] sequence, and is regulated by the m^6^A methyltransferase complex, and the two demethylases namely ALKBH5 and FTO^14^. Both demethylases have been emerged as potential promoters of cancer, acting at both in the nucleus and the cytoplasm^15^. They both regulate either mRNA stability or translation. In particular, mouse ALKBH5 was shown to drive ATF4 mRNA translation in amino acids-deprived mouse embryonic fibroblasts (MEF)^11^. The underlying mechanism involves cytoplasmic demethylation of a specific and conserved m^6^A (A225) located at uORF2 of ATF4 mRNA. This demethylation event forces the ribosomes to bypass the translation initiation codon of uORF2, allowing the selection of the downstream main translation initiation codon for ATF4 translation. Mouse FTO was similarly shown to promote ATF4 expression in amino acids-deprived MEFs^11^. However, whether ALKBH5 or FTO are required for ATF4 mRNA translation that occurs in chemotherapeutic-treated cancer cells to promote resistance, is still unknown.

We have previously shown that treatment of the hepatocarcinoma Hep3B cell line with the chemotherapeutic drug, sorafenib, induces an ER stress characterised by activation of phosphorylation of eIF2α and downstream induction of ATF4 mRNA translation, which occurs at the initiation step^16^. We further showed that SOR-induced ATF4 mRNA translation initiation also involves the activity of the RNA helicase DDX3 through a mechanism requiring interaction with both the translation initiation eIF4F factor and polyribosomes^10^. Preventing this SOR-induced ATF4 expression either directly via sh/siRNAs^16^ or indirectly by targeting its translational activator DDX3^10^, significantly sensitizes Hep3B to treatment. Here, we report that ALKBH5 and FTO are novel polyribosomes-associated proteins that drive ATF4 mRNA translation in the Hep3B treated with the chemotherapeutic sorafenib (SOR), promoting resistance.

## Materials and Methods

### Cell lines and Culture cell

Hepatocarcinoma Hep3B cells were previously described^10,16^. Hep3B and derivatives were maintained in Dulbecco’s modified Eagles’ medium (DMEM) (Wisent) supplemented with 5% heat-inactivated fetal bovine serum (FBS; Wisent), penicillin and streptomycin at 37 °C in 5% CO_2_.

### Establishment of stable depleted cell line, mall-interfering RNA (siRNA) experiments

ShRNA-mediated depletion of ALKBH5 or FTO was obtained using lentiviral shRNA. ALKBH5 shRNA was generated by ligation of oligonucleotides into the *Age*I and *Eco*RI restriction sites of pLKO.1 (Addgene plasmid #8453). Lentiviral-shRNA particles were generated by transfecting HEK 293T cells with 12 μg of pLJM1 vector containing the shRNA, 6 μg of psPAX2 packaging plasmid (Addgene plasmid #12260), and 2 μg pMD2.G envelope plasmid (Addgene plasmid #12259). Medium was changed 16 h after transfection and lentiviral particles were harvested 24 h later. Viral supernatant was filtered through 0.45 μm filters and supplemented with 8 μg/ml polybrene (Sigma). The supernatant was added to the cells for 24 h before the start of puromycin selection.

siRNAs were obtained from Dharmacon (Lafayette, CO). siRNA transfections were performed essentially using Hiperfect reagent (Qiagen) following the manufacturer’s protocol. Twenty-four hours before transfection, cells were trypsinized and plated in a 6-well plate at a density allowing to reach 50–60% confluence after twenty-four hours. Annealed duplexes were used at a final concentration of 10 nM. Forty-eight hours posttransfection, cells were treated with the same siRNA (5 nM) for an additional forty-eight hours before treatment.

The sequences of shRNA/siRNA used are:

siALKBH5 sense sequence: 5’ - GCU GCA AGU UCC AGU UCA A -3’
shALKBH5 sense sequence: 5’-CGG CAG AGT TGT TCA GGT T-3’
shFTO sense sequence: 5’ GCTGAGGCAGTTTTGGTTTCA-3’

### Stable GFP Transfection

mEGFP and mEGFP-ALKBH5 sequences were first ligated into the *Age*I and *PST*I restriction sites of a pLJM1 plasmid (Addgene). Lentiviral particles were generated by transfecting HEK 293T cells with 12 μg of pLJM1-mEGFP or pLJM1ALKBH5 plasmids with 6 μg of psPAX2 packaging plasmid and 2 μg of pMD2.G envelope plasmid (Addgene). The medium was replaced 16h after transfection, lentiviral particles were harvested 24h later, filtered through 0.45 μm filters, and supplemented with 8 μg/mL polybrene (Sigma). The collected viruses were added to Hep3B cells for 24h before Hep3B stably expressing mEGFP or mEGFP-ALKBH5 were then selected via puromycin resistance selection.

### Drugs treatment

Sorafenib was purchased from Selleck Chemicals. Sorafenib was dissolved in DMSO as a 10 mM stock solution, aliquoted and stored at −80 °C. MTT reagent was purchased from Sigma and dissolved in PBS 1X as a 5mg/ml solution. For drug treatment, cells were plated to reach a confluency of ∼80–90% on the day of the treatment. The media was changed with fresh media 2 hours before treatment.

### Antibodies

Anti-DDX3 and anti-tubulin antibodies were purchased from Abcam. Anti-ATF4, anti-ALKBH5, anti-FTO and anti-m^6^A antibodies were obtained from Proteintech. RPL22 et RPS14 (from Santa Cruz Biotech) were kindly provided by Dr Tom Moss (Université Laval). anti-FMRP antibodies were previously described^17^.

### Poly(ribo)some profiling and analyses of polysomal-associated protein and mRNAs

To prepare polysomes, we used the same protocol we previously described^18^. Briefly, cells grown in 100-mm tissue culture (∼ 80–90% confluence) were treated with sorafenib, then lysed with 1ml of polysomal buffer (20 mM Tris, pH 7.4, 150 mM NaCl, 1.25 mM MgCl_2_, 5 U/ml RNase inhibitor [Invitrogen], protease inhibitor cocktail [Complete; Roche], 1 mM DTT, and Nonidet P-40 at a final concentration of 1%. Extracts were clarified by centrifugation (12,000 *g*, 20min, 4°C) and the resulting cytoplasmic extracts were loaded on 15–55% (w/v) linear sucrose gradient for sedimentation by ultracentrifugation. RNA-proteins complexes of individual isolated fractions were ethanol precipitated. For protein analysis, proteins were resuspended in SDS-PAGE sample buffer. For RNA analysis, precipitated complexes were resuspended and fractions corresponding to monosomes, light and heavy polysomes were pooled and processed for RNA extraction and analyzed by RT-qPCR as described below.

### Biotinylated ATF4 RNA Reporter

In vitro transcribed and biotinylated ATF4 reporter RNA was produced as follows. Briefly, PCR-amplified DNA templates encoding 5’UTR of ATF4 mRNA were used for in vitro transcription of ATF4 reporter mRNA using HiScribe T7 High Yield RNA Synthesis Kit (NEB) according to the manufacturer’s protocols. The transcription products were purified with Monarch RNA Cleanup Kit (NEB) followed by polyadenylation using Poly(U) Polymerase (NEB) with Bio-16-UTP (Invitrogen) and purification with Monarch RNA Cleanup Kit (NEB). RNAs were then used for MeRIP assays.

### Methylated RNA immunoprecipitation (MeRIP) assay

For the MeRIP assay, total RNA was isolated from the Hep3B cells using Trizol reagent (Life Technology). Then, RNA was incubated overnight with ProteinA Sepharose beads coupled with anti-m^6^A antibody (Proteintech) or anti-IgG antibody in 500μl of IP buffer (recipe) at 4°C. After washes, the beads were incubated with proteinase K buffer (10 mM Tris-HCl (pH 7.4); 150 mM NaCl; 0.1% NP-40 substitute; 1 mM DDT, 8U/ml RNase inhibitor (Invitrogen), 2mg/ml Proteinase K and 0.1% SDS) at 55°C for 30min. Immunoprecipitated RNA was extracted, and the expression of related genes was detected via RT-qPCR.

For MeRIP assay using the biotinylated ATF4 RNA reporter, untreated and SOR-treated Hep3B were lysed in a lysis buffer (50 mM Tris-HCl (pH 7.4); 150 mM NaCl; 1mM MgCl2; 0.5% NP-40 substitute; 0.25 mM PMSF; 1 mM DDT, 5U/ml RNase inhibitor). After centrifugation, the supernatants were incubated with 5 µg of the biotinylated reporter overnight at 37°C, followed by incubation with proteinA beads coupled with anti-m^6^A antibody (Proteintech) or anti-IgG antibody at 4°C for 4h. After incubation with proteinase K buffer (10 mM Tris-HCl (pH 7.4); 150 mM NaCl; 0.1% NP-40 substitute; 1 mM DDT, 8U/ml RNase inhibitor, 2mg/ml Proteinase K and 0.1% SDS) at 55°C for 30min, methylated RNAs were extracted and then incubated with streptavidin-agarose beads (Thermo Scientific™) during 2h at 4°C. Following extraction, the methylated reporter was analyzed by RT-pPCR.

### DNA transfection and Luciferase reporter assay

Hep3B were transfected with a mixture of pRL-TK-Renilla, and the p5′UTR-ATF4-FLuc (WT ATF4-FLuc) or p5′UTR-ATF4 A235G-FLuc (mut ATF4-FLuc) plasmids using a Transfection reagent kit (Qiagen) as we previously described^10^. p5′UTR-ATF4 A235G-FLuc reporter mutant was produced by NorClone (London, ON, Canada). Forty-eight hours later, cells were treated with Thapsigargin (100nM, 4h) and cells were resuspended in passive lysis buffer (Promega). Firefly and *Renilla* luciferase activities were measured using a dual-luciferase reporter assay system (#E1960; Promega) according to the manufacturer’s instructions. Relative values of firefly luciferase activities were normalized to *Renilla* luciferase control.

### Quantitative Real-time PCR analysis

Total RNA was extracted with the Trizol reagent (Life Technology) and polyribosomal RNA was prepared by phenol-chloroform extraction. RNA was then reverse transcribed using the Quantitect Reverse Transcriptase kit (Qiagen). Real-time PCR reactions were prepared using the Luna® Universal qPCR Master Mix (NEB). Reactions were run and data analyzed using the QuantStudio^TM^ 7 Flex Real-time PCR system (Applied Biosystems). Data were calculated using the 2^-△△Ct^ method.

The primer sequences are:

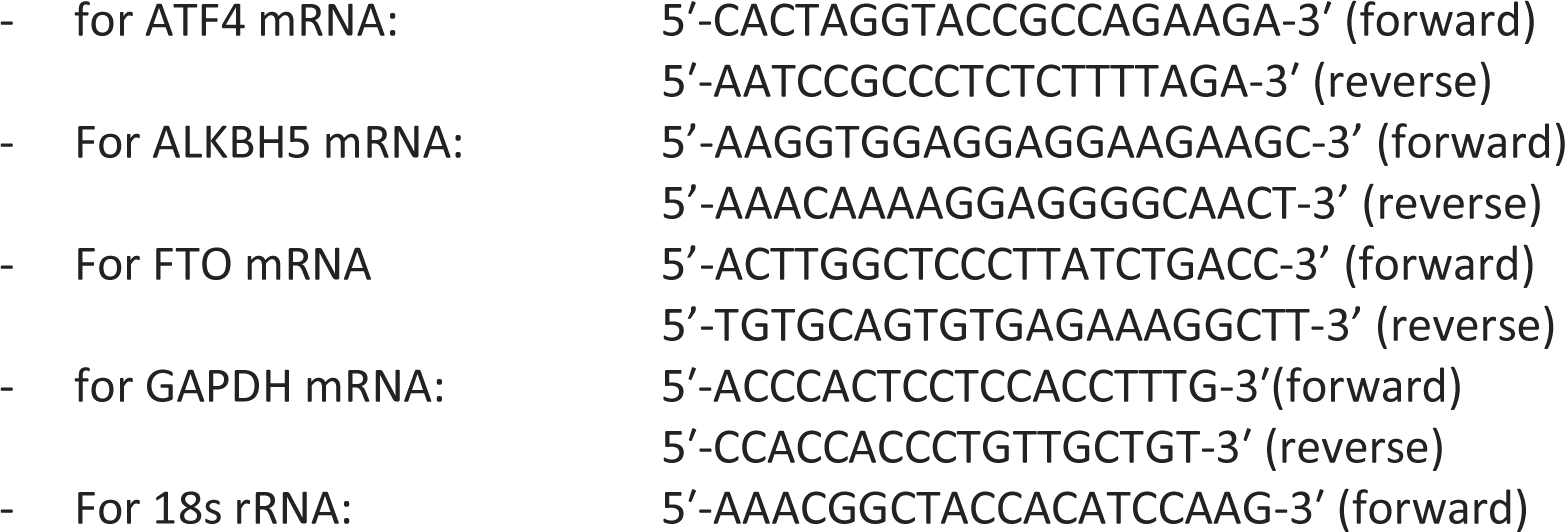
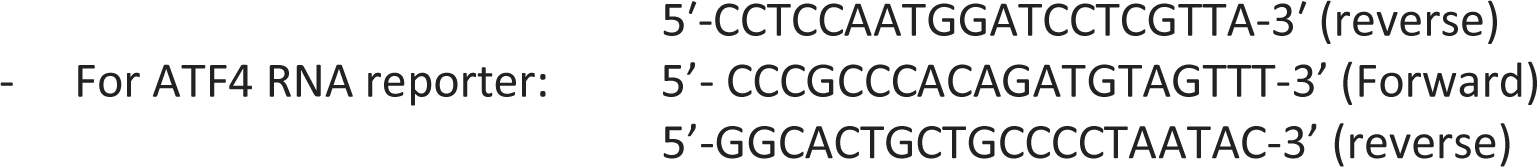

### GST Pull-down and GFP-Trap Assays

For GST pull-down, untreated or SOR-treated (10µM, 2h) Hep3B were lysed in extract in the following lysis buffer: 50 mM Tris-HCl (pH 7.4); 150 mM NaCl; 0.5% NP-40 substitute; 0.25 mM PMSF, and protease inhibitors. After clarification, cell lysates were incubated with purified GST and GST-DXX3 immobilized on glutathione agarose beads for 1h at 37°C. Beads were washed and boiled at 95 °C in 1x sodium dodecyl sulfate (SDS) loading buffer for 10 min and eluted proteins were analyzed by Western Blot. When indicated, the beads were incubated for 15 min at room temperature with RNase A (10 μg/mL) before processing.

For GFP-Trap assay, Hep3B stably expressing GFP or GFP-ALKBH5 were lysed in 1 mL lysis buffer (50 mM Tris-HCl pH 7.4, 150 mM NaCl, 1 mM MgCl2; 0.5% NP-40 substitute, 0.25 mM phenylmethylsulfonyl fluoride, 0.1 mM dithiothreitol, and a protease inhibitor cocktail added immediately before processing). After clarification, the lysates were incubated for 2h at 4 °C with GFP-Trap-agarose beads (Proteintech). The beads were then washed in lysis buffer and boiled at 95 °C in Laemmli buffer for 10 min to eluate the bound proteins. The eluted proteins were then analysed by Western blot. When indicated, the beads were incubated for 15 min at room temperature with RNase A (10 μg/mL) before processing.

### MTT assay

For MTT assays, cells were seeded at 10 000 cells/well in 96-well plates. Cells were cultured overnight and then treated with 10 μM sorafenib for 24h. At the end of the treatment, cells were washed with PBS and complete media was added. At the end of treatment, cells were washed with PBS and 3-(4,5-dimethyl-2-thiazolyl)2,5-diphenyl-2-H-tetrazolium bromide (MTT, Sigma-Aldrich) solution (final concentration: 0.5 mg/mL) was added to each well. Following 2h of incubation, the reaction was stopped with 150 μL DMSO and the cells were shaken for 5 min at room temperature. The absorbance of each well was measured at 560 nm and the viability of cells was calculated.

### Clonogenic assay

After treatment with sorafenib (10μM, 24h), cells were trypsinized, counted, replated in 6-well plates at 1000 cells/well, and incubated for 8–10 days. Cells are then washed with PBS and stained (0.1% (w/v) crystal violet in a 0.0037% (v/v) formaldehyde solution in PBS). Isolated colonies were counted to determine cell viability.

**Immunofluorescence** (supplementary data)

For immunofluorescence experiments, all fixation, permeabilisation and staining procedures were performed according to the previously described protocol^10^. Briefly, after treatment, cells were fixed with 3,7% paraformaldehyde and permeabilized with methanol. The sample was blocked with 1% BSA and incubated with primary antibodies diluted in PBS containing 1% BSA and 0,1% Tween-20. Cells were then incubated with secondary antibodies coupled to Alexa Fluor 488 or 594 (Life Technology). Immunostainings were visualized using an LSM 900 laser scanning confocal microscope (Zeiss) controlled with ZENsoftware for image acquisition and analysis.

## Results

### Depletion of either ALKBH5 or FTO downregulates SOR-induced expression of ATF4 mRNA

It was previously reported that mouse ALKBH5 drives ATF4 mRNA translation in amino acids-deprived mouse embryonic fibroblasts (MEF)^11^. The underlying mechanism involves demethylation of a specific m^6^A225 located at uORF2 of ATF4 mRNA, which allows the ribosomes to select the main translation initiation codon for translation^11^. Mouse FTO was similarly shown to promote ATF4 expression in amino acids-deprived MEFs^11^, potentially through a similar mechanism, though this was not investigated. Because this m^6^A225 site is conserved in mammals (m^6^A235) (Fig. 1A), we postulated that a similar epitranscriptomic mechanism involving the activity of ALKBH5 and FTO may contribute to the induction of ATF4 mRNA expression in mammalian cells, and thus may trigger this translation in cancer cells treated with chemotherapeutics. To test this, we used the Hep3B treated with SOR that we have previously shown to induce translation of ATF4 mRNA via both phosphorylation of eIF2α^16^ and DDX3 activity^10^. To assess the effects of depleting the demethylases ALKBH5 and FTO in regulating the expression of SOR-induced ATF4 mRNA, we first generated, through a lentivirus system, Hep3B-stably expressing shRNAs that target either the coding sequence or 3’UTR of ALKBH5- and FTO-mRNAs, depleting the corresponding proteins, as validated by western blot analysis using specific antibodies (Fig. 1B-C). As shown in figure 1B-C and Supplementary data 1, depletion of either ALKBH5 or FTO significantly reduces SOR-induced ATF4 expression, indicating a role of either protein in ATF4 mRNA expression. This reduction of ATF4 mRNA expression occurring during the first two hours of SOR treatment, indicates that it is likely due to its altered induction. Control western blot experiments show that depletion of ALKBH5 or FTO has no major effect on SOR-induced eIF2α phosphorylation (Fig. 1B-C), and RT-qPCR analyses (Fig. 1D) showed that depleting ALKBH5 or FTO does not affect the level of ATF4 mRNA, indicating that ATF4 mRNA expression mediated by ALKBH5 and FTO occurs posttranscriptionally and independently of eIF2α phosphorylation. Together, these results indicate that both ALKBH5 and FTO contribute to the induction of expression of ATF4 mRNA during ER stress triggered by SOR treatment of Hep3B.

**Figure 1:**
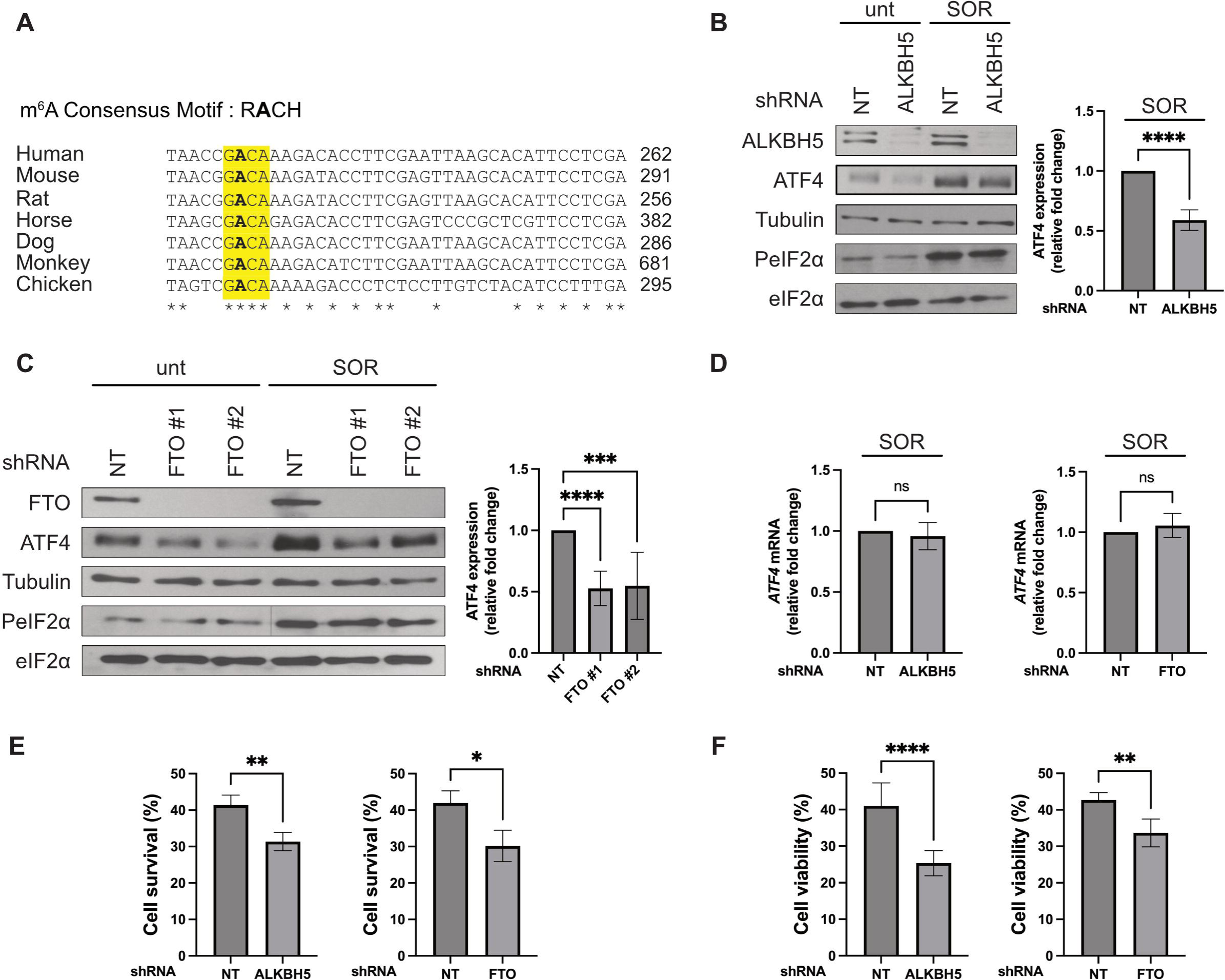
The RNA demethylases ALKBH5 and FTO are required for SOR-induced ATF4 expression. (A) Conserved consensus m^6^A site in the 5’UTR of ATF4 mRNA. R = A or G; H = A, C or U. * corresponds to the conserved nucleotides including the methylated adenosine in bold. (B-D) Hep3B were treated with 10µM SOR for two hours. (B) Left panels: Cells were harvested, lysed and protein extracts were analyzed by western blot for the expression of ALKBH5, ATF4, PeIF2α, the pan-eIF2α, and tubulin (Tub; loading control) using the corresponding antibodies. Right panel: The expression level of ATF4 was estimated by densitometry quantification of the film signal using Image Studio™ Lite Software and standardized against total tubulin. ****P ≤ 0.0001 (Student’s t-test). (C) Left panels: Cells were harvested, lysed and protein extracts were analyzed by western blot for the expression of the indicated proteins using the corresponding antibodies. Tubulin serves as a loading control. Right panel: The expression level of ATF4 was estimated by densitometry quantification of the film signal using Image Studio™ Lite Software and standardized against total tubulin. ***P ≤ 0.001; ****P ≤ 0.0001 (Student’s t-test). (D) Total RNA was isolated from harvested Hep3B and the level of ATF4 mRNA relative to GAPDH mRNA was quantified by real-time q(RT)-PCR using the ΔΔCt method. The presented results are the mean of at least triplicate measurements, with error bars corresponding to the S.D. (E-F) Hep3B were treated with 10µM SOR for twenty-four hours. (E) Clonogenic survival assays. After treatment, cells were trypsinized, counted and seeded in the absence of drug, and incubated for 10 days. Populations > 20 cells were counted as one surviving colony. Data were calculated as the percentage of surviving colonies relative to the number found in plates corresponding to mock-depleted cells plates. Data shown are representative of at least 3 separate experiments and values are given as mean ± SD. (Student’s t-test). **P ≤ 0.01; *P ≤ 0.05. (F) Cell viability, assessed by MTT assay, shows the viability of Hep3B cells stably expressing shALKBH5 or shFTO after exposure to SOR for 24h. Data shown are representative of at least 3 separate experiments and values are given as mean ± SD. (Student’s t-test). **P ≤ 0.01; ****P ≤ 0.0001

We have previously shown that eliminating SOR-induced ATF4 expression either directly by depleting ATF4 mRNA with specific si/shRNA or indirectly by depleting its upstream DDX3 regulator reduced Hep3B survival^10,16^. Our results implicating ALKBH5 and FTO in SOR-induced ATF4 expression prompted us to test if the demethylases contribute to SOR resistance. Hep3B stably expressing either shALKBH5 or shFTO, were treated with SOR and then collected and analyzed for survival by both clonogenic and MTT assays. Depletion of ALKBH5 and FTO significantly reduces the survival of SOR-treated Hep3B (Fig. 1E-F) indicating that both ALKBH5 and FTO promote cell resistance to SOR. Collectively, these results indicate that ALKBH5 and FTO drive SOR resistance, possibly by promoting the induction of ATF4 expression.

### ALKBH5 and FTO are polysome-associated proteins that promote the loading of ATF4 mRNA into translating ribosomes upon ER stress

Both ALKBH5 and FTO have been described as nuclear proteins, though they have been implicated in cytoplasmic functions^14^. We thus tested if the ALKBH5 and FTO-mediated regulation of ATF4 expression during SOR treatment reflects a specific localisation of the proteins. Due to the lack of anti-ALKBH5 suitable for immunofluorescence experiments, we used our Hep3B stably expressing GFP-ALKBH5. As expected, GFP-ALKBH5 distributes mainly in the nucleus of Hep3B (Supplementary data 2). We have previously shown that SOR treatment induces the formation of cytoplasmic stress granules (SG) that contain DDX3^16^. We did not detect GFP-ALKBH5 in SOR-induced SG containing DDX3, which is consistent with previous data failing the detection of the protein in SG induced by arsenite in U2OS cells, using both localisation^19^ and proteomics studies^20^. FTO was detected in the nucleus in mammalian cells such as U2OS, while no protein was detected in SG that form in Arsenite-treated cells^19^. Similarly, we found FTO mainly nuclear in Hep3B (Supplementary data 2). In response to SOR treatment, FTO was not detected in DDX3 positive-SG (Supplementary data 2). Collectively, these results excluded an association of ALKBH5 and FTO with SOR-induced SG, which is consistent with their role in promoting the expression of ATF4 mRNA, potentially at the translational level.

The induction of ATF4 mRNA expression that occurs during treatment with SOR is mainly translational^10,16^. Our data showing that depleting either ALKBH5 or FTO prevents SOR-induced ATF4 expression suggested a possible role of either protein in ATF4 mRNA translation. To investigate this possibility, we first assessed ALKBH5 and FTO association with the translational machinery in untreated and SOR-treated Hep3B by polysome profiling analysis. Cytoplasmic extracts of untreated or SOR-treated Hep3B were processed through sucrose density gradients followed by separation of translation initiation complexes and translating polysomes fractions (Fig. 2A). The polysome profiles are validated by western blot analysis of cell extracts of isolated fractions using anti-ribosomal RPL22, -RPS14 and -FMRP antibodies (Fig. 2, bottom panel). In physiological cell growth condition, we found that both ALKBH5 and FTO distributed through the gradient (Fig. 2A, left panel), suggesting that the demethylases associate with both translation initiation complexes, sedimenting at the top of the gradients, and with translating polysomes sedimenting at the middle and bottom of the gradient, identifying the proteins as polysomes-associated proteins. As we have previously described^16^, treatment of Hep3B with SOR induces a significant collapse of polysomes (Fig. 2A, right top panel), indicating a general inhibition of translation initiation. Despite the loss of the majority of polysomes in SOR-treated Hep3B, ALKBH5 and FTO are still detected, albeit slightly in fractions corresponding to the residual translating polysomes (Fig. 2, right bottom panel), further supporting the possibility that ALKBH5 and FTO promote the induction of ATF4 mRNA translation upon SOR treatment.

**Figure 2.**
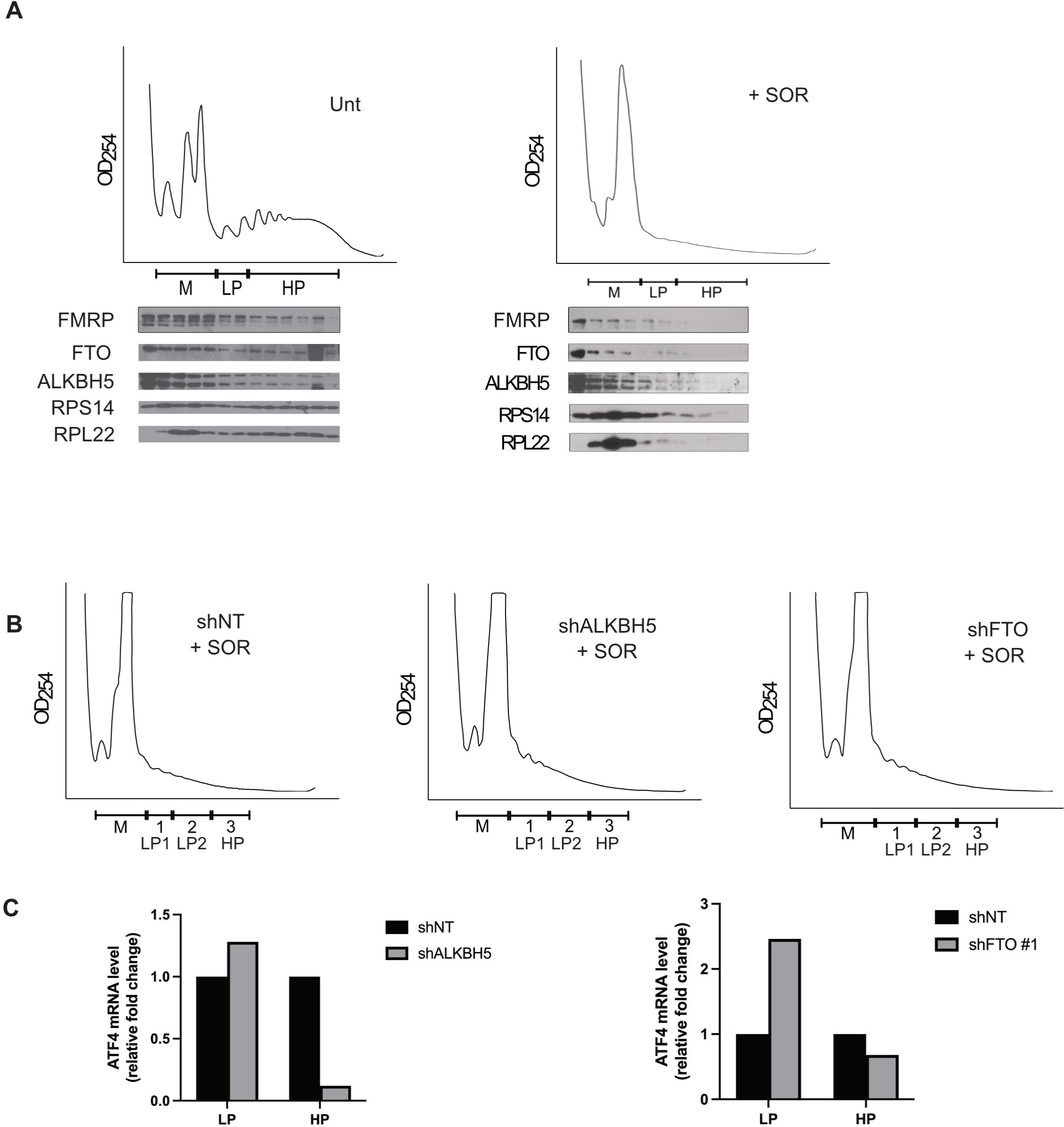
Both ALKBH5 and FTO are required for the association of ATF4 mRNA with polysomes in SOR-treated Hep3B. (A) Analysis of ALKBH5 and FTO association with polysomes. Top panels: cytoplasmic extracts prepared from either untreated or SOR-treated Hep3B were fractionated through 15–55% (w/v) sucrose density gradients and their polysomes profiles were monitored by measuring the OD254. LP: light polysomes. HP: heavy polysomes. Bottom panels: Western blot analysis of the collected fractions for the distribution of FMRP, ALKBH5 and FTO using specific antibodies. RPS14 and RPL22 ribosomal proteins are used as controls for the integrity of the polysome profiles. The results are representative of two independent experiments. (B-C) Analysis of the ATF4 mRNA association with polysomes in the absence of RNA demethylases ALKBH5 and FTO. Hep3B stably expressing shRNAs against ALKBH5 or FTO or a non-specific shRNA (NT) control were treated with SOR (10µM, 2h) as above. Cytoplasmic extracts were fractionated through 15%-55% sucrose gradients and their polysomes profiles were recorded as above. (B) Polysome profile of SOR-treated Hep3B shNT, shALKBH5 and shFTO. (C) RNA content was isolated from pooled LP and HP fractions and associated ATF4 mRNA was quantified by RT-qPCR using the ΔΔCt method. ATF4 mRNA levels were normalized against 18S ribosomal RNA and expressed as indicated. The results are representative of two independent experiments.

We have previously shown that the association of a fraction of ATF4 mRNA with polysomes in SOR treatment of Hep3B and its translation are significantly prevented upon depletion of DDX3, downregulating its translation^10^. We then investigated if the depletion of ALKBH5 or FTO similarly affects the association of ATF4 mRNA with polysomes in SOR-treated Hep3B. In control experiments, depletion of ALKBH5 and FTO had no effect on the loss of polysomes induced by the treatment with SOR (Fig. 2B). Using RT-qPCR analysis of polysomes-associated mRNAs, we observed that depletion of ALKBH5 or FTO resulted in a significant decrease of the association of ATF4 mRNAs with polysomes (Fig. 2C, HP fraction), indicating that ATF4 mRNA translation is inhibited. Together, these results indicate that ALKBH5 and FTO promote SOR-induced ATF4 mRNA translation, potentially at the initiation step involving the demethylation of the 5’UTR of ATF4 mRNA.

### ALKBH5-mediated demethylation ATF4 mRNA negatively regulates its SOR-induced translation

High-throughput sequencing of modified RNAs using approaches such as the methylated RNA immunoprecipitation (MeRIP) showed that m^6^A modifications are mainly enriched in 5’ UTRs, 3’ UTRs and near stop codons^21,22^, and that the level of modified RNAs increases in response to stress such as oxidative stress^19^ and heat shock^23^. In agreement, we found that treatment of Hep3B with SOR significantly increases the overall level of m^6^A of mRNAs as assessed in m^6^A dot blotting of Hep3B-purified poly(A) mRNAs using anti-m^6^A antibodies (data not shown). We then performed MeRIP combined with RT-qPCR analyses to determine the m^6^A level of ATF4 mRNA in SOR-treated Hep3B. We observed ∼2-fold increases in methylation of ATF4 mRNA in SOR-treated Hep3B as compared to untreated Hep3B (Fig. 3A), indicating that SOR treatment induces a global methylation level of ATF4 mRNA, probably occurring at the multiple m^6^A sites found in its coding sequence, as previously reported by MeRIP-seq in HEK293 or U2OS during both oxidative stress and heat shock^19,23–25^. Similarly, several sites of ATF4 mRNA were shown to be methylated in amino acid-starved mouse embryonic fibroblasts, however, the conserved m^6^A225 located at the 5’UTR of ATF4 mRNA exhibited reduced methylation, an event required for its translation^23^. To investigate if this specific demethylation similarly occurs in SOR-treated Hep3B, we performed a modified MeRIP using an *in vitro* transcribed and biotinylated RNA reporter (Fig. 3B-C). Briefly, the 5’UTR of human ATF4 mRNA was biotinylated *in vitro* and incubated without (mock) and with similar amounts of extracts prepared from either untreated- or SOR-treated Hep3B to allow methylation, then subjected to methylated RNA immunoprecipitation (MeRIP) using m^6^A antibodies, and as control with IgG. IPed RNAs are then incubated with streptavidin-agarose beads to purify the biotinylated reporter RNA, which is quantified by RT-qPCR using oligos specific to the 5’end of ATF4 mRNA. The results show that incubation of the reporter RNA with Hep3B extracts efficiently induces its methylation, which was however significantly reduced upon SOR treatment (Fig. 3C). Thus, while incubation of the reporter ATF4 RNA with extracts of untreated Hep3B induces its methylation most likely at the conserved A235 (Fig. 1A), the lack of methylation of the reporter RNA in the extract of SOR-treated Hep3B supports our assumption that this treatment antagonizes m^6^A235 modification, potentially by activating a methylation antagonizing pathway, which may involve demethylases. Consistently, methylation of the reporter ATF4 mRNA was significantly enhanced upon incubation with extracts prepared from ALKBH5-depleted Hep3B and treated with SOR (Fig. 3D), indicating that alteration of the methylation of the reporter mRNA in SOR-treated Hep3B occurs through a mechanism involving the activation of an ALKBH5 pathway. Control experiments (Fig. 3D) showed that depletion of ALKBH5 does not affect the methylation status of the reporter mRNA incubated with the extracts from untreated Hep3B, indicating that ALKBH5 acts specifically upon SOR treatment for demethylation. Although our experiments do not assess directly the demethylation of the reporter, it supports the possibility that ALKBH5 interferes with its methylation, potentially at A235, enhancing its translation during stress.

**Figure 3.**
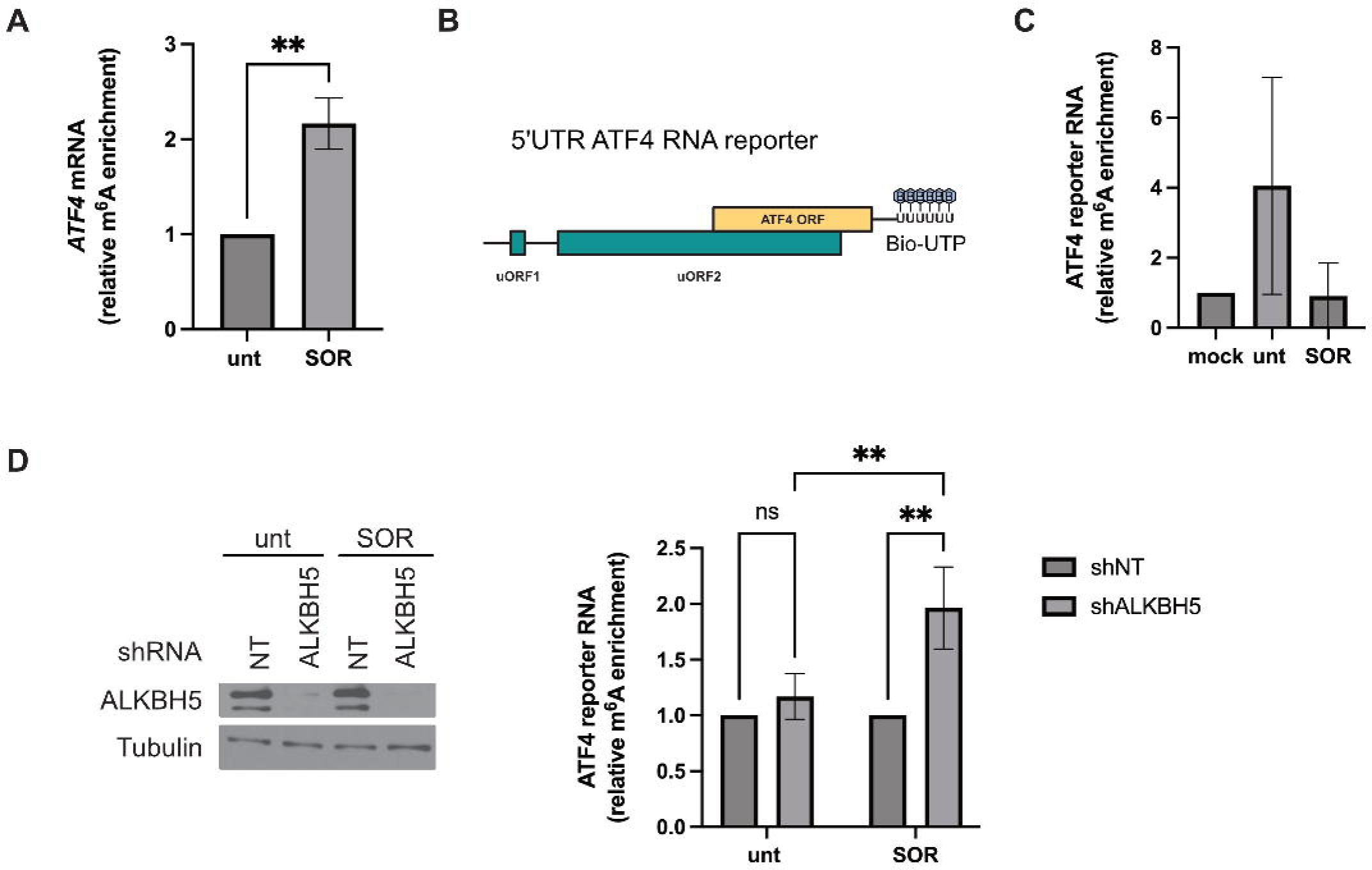
Role of ALKBH5 in ATF4 mRNA methylation level. (A) MeRIP-qPCR analysis of m^6^A level of ATF4 mRNA incubated with protein extracts prepared from untreated and SOR-treated Hep3B. m^6^A methylated RNAs were isolated and ATF4 mRNA was quantified by RT-qPCR. The amounts of m^6^A ATF4 mRNA were normalized against IgG precipitate and then expressed relative to untreated condition. **P ≤ 0.01. (B) Schematic representation of the 5’UTR ATF4 RNA reporter. (C) MeRIP-qPCR analysis of m^6^A level of 5’UTR ATF4 RNA reporter incubated with protein extracts prepared from untreated and SOR-treated Hep3B. m^6^A methylated ATF4 reporter RNAs were isolated and quantified by RT-qPCR. The amounts of m^6^A ATF4 reporter RNA were normalized against IgG precipitate and then expressed relative to mock condition. (D) MeRIP-qPCR analysis of m^6^A level of 5’UTR ATF4 RNA reporter incubated with proteins extracts from Hep3B stably expressing either a control shRNA (shNT) or shALKBH5 and treated with SOR (10µM, 2h). Depletion of ALKBH5 is validated by western blot using specific antibodies as described in figure 1 (left panels). m^6^A methylated ATF4 reporter RNAs were isolated and quantified by RT-qPCR (right graphs). The amounts of m^6^A ATF4 reporter RNA were normalized against IgG precipitate and then expressed relative to shNT condition. Data are representative of 3 separate experiments and values are given as mean ± SD. **P ≤ 0.01.

To assess the assumption that ALKBH5-mediated demethylation of A235 of ATF4 mRNA regulates its translation, we devised the standard luciferase translational reporter assay that is widely used to assess translation of uORFs-harboring mRNAs^10^. Briefy, mock- and ALKBH5-depleted Hep3B are co-transfected with either a luciferase expressing vector containing the human ATF4 5′UTR fused to Firefy luciferase (FLuc) gene, or its mutated version (mutATF4-FLuc) by introducing an A235G mutation (Fig. 4A), together with the control plasmid expressing Renilla luciferase (RLuc). Cells are then treated with the drug to induce translation of FLuc whose activity is measured in the cell extracts and expressed relative to RLuc. As we previously described^10^, SOR treatment does not induce efficient translation of the reporter in this assay, as compared to treatment with the canonical ER stress inducer Thapsigargin (data not shown), precluding accurate and statistical quantification of FLuc level in SOR-treated cells. Thus, for these experiments, we used Thapsigargin, which we have shown to efficiently induce translation of both endogenous ATF4 and the F-luc reporter^10^. As expected, our control experiments showed that thapsigargin treatment of either Hep3B (Fig. 4B) or mock-depleted Hep3B (Fig. 4C) significantly induced translation of both the WT- and mut-ATF4-FLuc. Thapsigargine-induced translation of the WT ATF4-Fluc, but not of the control mut ATF4-FLuc reporter, was prevented in ALKBH5-depleted Hep3B (Fig. 4D), supporting that ALAKBH5 drives ATF4 mRNA translation, at least in part by demethylating A235, potentially involving interaction with ATF4 mRNA partners acting as m^6^A readers.

**Figure 4.**
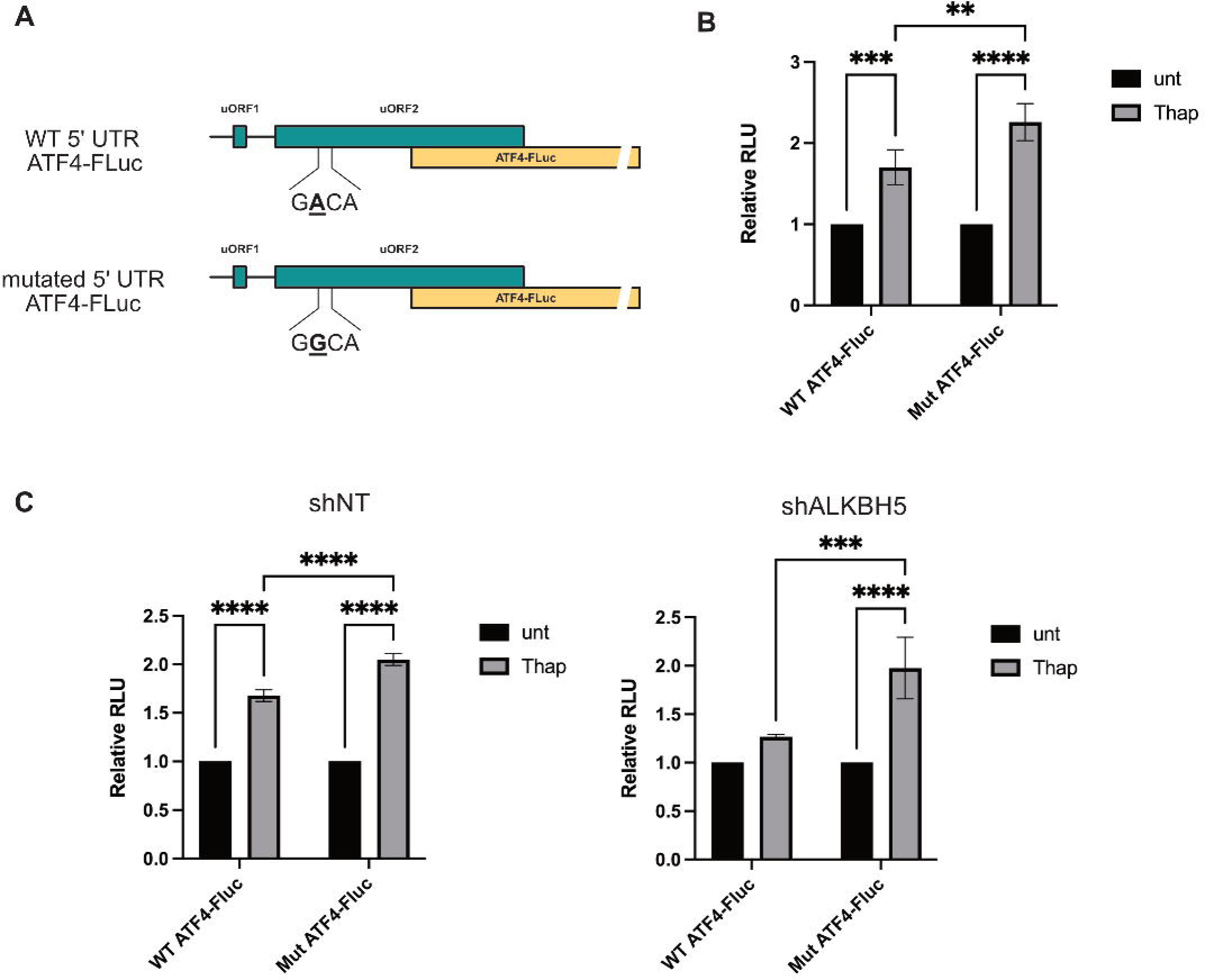
(A) Schematic representation of the 5’UTR ATF4 Luciferase reporters (WT- and mut-ATF4 FLuc) consisting of the human WT- and A235G (mut)-ATF4 5′-UTR fused to Firefly luciferase (FLuc) gene. (B) Hep3B cells are co-transfected with either Fluc expressing vector, and the control plasmid expressing *Renilla* luciferase (RLuc). Cells are then treated with Thapsigargin (thap) to induce translation of FLuc whose activity is measured in the cell extracts and expressed relative to RLuc. The relative values of firefly luciferase were shown as the average of three biological replicates. Error bars correspond to the S.D. **P ≤ 0.01, ***P ≤ 0.001, ****P ≤ 0.0001. (C) Hep3B stably expressing either a control shRNA (shNT) or shALKBH5 are co-transfected with a luciferase expressing vector containing the human ATF4 5′-UTR fused to FLuc gene, and the control plasmid expressing RLuc. Cells are then treated with Thap to induce translation of FLuc whose activity is measured. The relative values of firefly luciferase were shown as the average of three biological replicates. Error bars correspond to the S.D. ***P ≤ 0.001, ****P ≤ 0.0001.

### Both ALKBH5 and FTO interact with DDX3

We have previously reported that SOR-induced ATF4 mRNA translation requires the activity of DDX3^10^, a previously identified ALKBH5 partner^26^. Our finding that ALKBH5 and FTO are similarly required for this ATF4 mRNA translation, suggested the possibility that interaction between these demethylases with DDX3 is required for SOR-induced ATF4 mRNA translation. We thus sought to assess if these interactions occur in Hep3B and test if SOR treatment triggers their regulation. We first investigated if DDX3 and ALKBH5 interact in Hep3B using GFP-Trap (Fig. 5A). In these experiments, we performed pull-downs on cell extracts prepared from a similar number of Hep3B cells expressing either GFP-ALKBH5 or the control GFP, treated or not with SOR. Bound DDX3 was then evaluated by western blot analysis, validating the association of DDX3 with GFP-ALKBH5 in Hep3B. We observed a similar amount of DDX3 associating with GFP-ALKBH5 between untreated- and SOR-treated Hep3B (Fig. 5A), indicating that SOR treatment does not affect their interaction. In control experiments, RNase treatment had not effect on the interaction between GFP-ALKBH5 and DDX3, indicating that DDX3-ALKBH5 interactions are RNA-independent, mediated by specific motifs. In reciprocal GST pull-down experiments, we confirmed the association between purified recombinant GST-DDX3 immobilized on glutathione beads and endogenous ALKBH5 present in Hep3B treated or not with SOR detected by western blot, while in control experiments no interaction between GST and ALKBH5 was observed (Fig. 5B). As expected, RNase treatment had no effect on GST-DDX3 interaction with ALKBH5, further indicating that the two proteins associate directly (Fig. 5B). Using similar GST pull-downs, we also found a novel interaction between recombinant GST-DDX3 and FTO (Fig. 5C). Together, our results show an interaction between DDX3 with both demethylases in SOR-treated cells, which may be required for ATF4 mRNA translation, though we did not demonstrate this.

**Figure 5.**
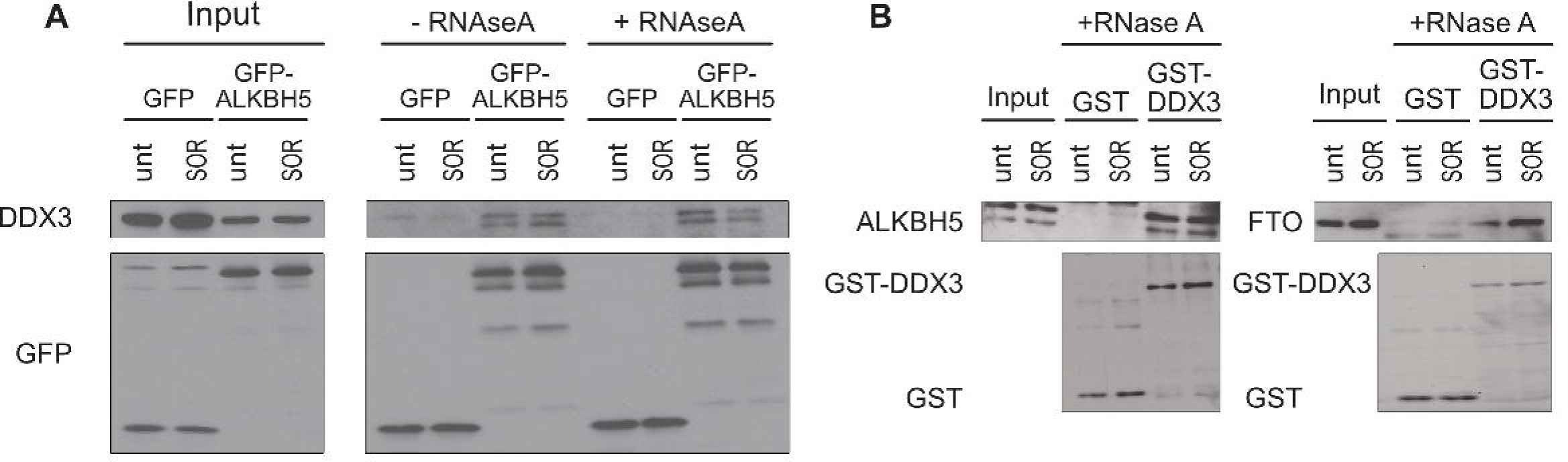
DDX3 interacts with both ALKBH5 and FTO. (A) GFP-Trap. Hep3B stably expressing GFP or GFP-ALKBH5 were treated with SOR (10 µM, 2h) or left untreated. After lysis, cell extracts were incubated with GFP-Trap-agarose beads for 2h at 4°C. Beads were then treated or not with RNAseA (10µg/mL, 15min at RT) and bound proteins were eluted and analyzed by Western blot using specific antibodies. (B) GST pull-down. Hep3B cells were treated with SOR (10 µM, 2h) or left untreated and the cell lysates were incubated with GST or GST-DDX3 immobilized on glutathione beads. Before elution, beads were treated or not with RNase A (10µg/mL, 15min at RT), and bound proteins were then analyzed by Western blot using specific antibodies.

## Discussion

In this study, we found that both RNA demethylases ALKBH5 and FTO promote ATF4 mRNA expression that is induced by SOR treatment. Our finding that either demethylase associates with polysomes and that their respective depletion abrogates the association of ATF4 mRNA with translating ribosomes in SOR-treated Hep3B supported the role of either demethylase in promoting the expression of ATF4 mRNA at the translational level. The findings that the level of methylation of the 5’UTR of ATF4 mRNA is increased upon depletion of ALKBH5, and that depletion of ALKBH5 abrogates the translation of the WT-but not the mut-ATF4 FLuc reporter, are consistent with the role of the protein in antagonizing m^6^A, supporting a conserved RNA editing mechanism required for stress-induced ATF4 mRNA translation. Finally, our data showing that ALKBH5 and FTO are required for Hep3B to resist SOR treatment, underscored the potential role of RNA demethylases in chemoresistance, possibly by allowing translation of ATF4 mRNA.

Two studies reported a role of ALKBH5 in stress-mediated ATF4 mRNA expression, either by promoting its induction at the onset of stress or sustaining its expression during prolonged stress. In Zhou et al. study, mouse ALKBH5 was reported to be required to demethylate A225 of ATF4 mRNA and drives its translation in MEFs during the first 2 hours of starvation^11^, indicating that ALKBH5, and possibly FTO, are recruited to the ATF4 mRNA at the early phase of stress inducing its demethylation and translation, though it remained unknown how this recruitment event occurs. Our data showing that 2 hours of SOR treatment of Hep3B is sufficient to induce ATF4 mRNA translation, while this induction is prevented by depleting either ALKBH5 or FTO, support the role of demethylases at the early phase of stress required for the induction of ATF4 mRNA translation. The study of Mukhopadhyay et al. supported however a possible alternative ALKBH5-based regulatory mechanism of ATF4 mRNA translation^12^. In this mechanism, ALKBH5 is required to sustain the expression of ATF4 mRNA that occurs during persistent ER stress that is induced by prolonged (over 8 hours) thapsigargin treatment of human HEK293T^12^. ALKBH5-mediated sustaining of ATF4 expression in thapsigargin-treated cells requires a significant increase in the expression level of ALKBH5^12^. We did not observed however any change in the expression of ALKBH5 during SOR treatment (Supplementary data 1B), even during prolonged treatment, which is consistent with our assumption that ALKBH5 is required for the induction step of ATF4 translation during drug treatment. This is also consistent with our finding that depletion of ALKBH5 significantly abrogates translation of the FLuc reporter that occurs during acute thapsigargin treatment (Fig. 4). Thus, the above studies together with ours support a translational role of ALKBH5, involving alternate stress- or cell type-specific mechanisms, which may also depend on its expression level, and binding partners.

How ALKBH5 is recruited to the ATF4 mRNA to induce its demethylation remains to be identified. We have previously shown that DDX3 is required for SOR-induced ATF4 mRNA translation, through a mechanism involving interaction with eIF4F, which is responsible for the recognition of the m^7^G cap structure at the 5’end of ATF4 mRNA during the initiation of its translation^10^. While no interaction between ALKBH5 and eIF4F was reported, Shah et al. also showed that DDX3 interacts with ALKBH5 in HEK293 cells^26^. Using both GFP-TRAP and GST pull-down, we confirmed an interaction between DDX3 and ALKBH5 in Hep3B cells (Fig. 5), raising the possibility that DDX3 may act as an m^6^A reader recruiting demethylases to ATF4 mRNA, inducing its demethylation and translation in SOR-treated Hep3B. In this case, DDX3 may serve a dual function in the promotion of the ATF4 mRNA translation initiation: DDX3 recruits the eIF4F cap-binding complex to the 5’cap of ATF4 mRNA allowing its recognition by the translation initiation machinery, while it recruits demethylases, which by demethylating the conserved A235 allows the translation machinery to translate the main ORF. Clearly, future studies are needed to investigate the proposed dual role of DDX3, involving interaction with ALKBH5, and possibly FTO demethylases. While Shah et al. study excluded interaction between DDX3 and FTO in HEK293T, our GST pull-downs revealed this interaction in Hep3B, potentially driving ATF4 mRNA translation, though we did not address this here. Additional studies are required to characterize the DDX3 interaction with RNA demethylases and define its functional role in promoting SOR-induced ATF4 mRNA translation, potentially by demethylating functional m^6^A.

Our studies using the reporter biotinylated ATF4 mRNA (Fig. 3) are consistent with Zhou et al. showing that the A225 in mouse ATF4 mRNA, corresponding to the A235 in its human orthologue, is the main ALKBH5 target site for demethylation to promote ATF4 mRNA translation. In addition to m^6^A, both mouse and human ATF4 mRNA can to be methylated at the first transcribed and conserved adenosine adjacent to the m^7^G cap, through a modification called *N*^6^,2’-O-dimethyladenosine (m^6^Am)^12,27^. However, whether demethylation of m^6^Am is required for the induction of ATF4 mRNA translation upon stress, remains unknown. Demethylation of m^6^Am is posed to be mediated mainly via FTO^28,29^, regulating the stability of target mRNAs, with minimal effect on their translation^27^. Our data showing that the reporter ATF4 mRNA, which is uncapped and lacks the first potential m^6^Am adenosine is readily methylated in Hep3B extracts indicated however that A335 is the main residue of ATF4 mRNA that is targeted by methylation/demethylation, regulating its stress-induced translation in our cell models. This is further supported by our luciferase reporter assays showing that translation of the capped ATF4-FLuc reporter harboring an A/G235 mutation is resistant to ALKBH5 depletion. Nevertheless, we do not exclude the contribution of m^6^Am in the process. Clearly, more studies including reporter assays using capped 5’UTR ATF4 mRNA and harboring an A/G235 mutation that prevents its m^6^A modification are required to determine the m^6^Am methylation status of the reporter in Hep3B extracts, while assessing if its potential demethylation in drug-treated cells involves the activity of FTO, potentially involving interactions DDX3.

We have previously shown that SOR treatment induced a dynamic repartition and localization of ATF4 mRNA^16^. We found that while a sub-fraction of ATF4 mRNA is associated with polysomes for translation, a significant fraction localizes into SG, potentially in a translation repressed form^16^. At this stage, we do not know if the association of ATF4 mRNA with SG and polysomes is mediated by its methylation and demethylation at A235, respectively. Nevertheless, this assumption is consistent with reports showing that mRNAs harboring high levels of m^6^A, but not those lacking m^6^A are enriched in SG^19^, which is also supported by data^19^ showing an enrichment of m^6^A signal in SG induced in arsenite treated-cells, further supporting the possibility that modified RNAs are selectively sorted to SG, while demethylated mRNAs may be preferentially translated. In any case, future studies validating the role of ALKBH5 and FTO in demethylating the ATF4 mRNA during SOR treatment, combined with experiments assessing the localisation of reporter ATF4 mRNAs in SOR-induced SG, should help understand the dynamic role of m^6^A in the repartition of ATF4 mRNA between SG and polysomes, regulating its translation.

### Supplementary data

Supplementary Figure 1. (A) Hep3B were treated with ALKBH5 or control (NT) siRNAs for ninety-six hours and then incubated with 10 μM SOR for two hours. Left panel: Cells were harvested, lysed and protein extracts were analyzed by western blot for the expression of ATF4, ALKBH5 and tubulin (Tub; loading control) using the corresponding antibodies. Right panel: The expression level of ATF4 was estimated by densitometry quantification of the film signal using Image Studio™ Lite Software and standardized against total tubulin. **P ≤ 0.01 (Student’s t-test). (B) Hep3B were treated with 10 μM SOR for the indicated times. Cells were collected, and their protein extracts were analyzed by western blot for the expression of ATF4, ALKBH5 and DDX3. Tubulin serves as a loading control.

Supplementary Figure 2. (A) Hep3B stably expressing either GFP-ALKBH5 or the control GFP were treated with 10µM SOR for two hours and then processed for immunofluorescence using anti-DDX3 antibodies (stress granule maker). Images were acquired by the LSM 900 confocal microscope. Scale bars (10 μm) are shown. (B) Hep3B were treated with 10µM SOR for two hours and then processed for immunofluorescence using anti-FTO and anti-FMRP antibodies. FMRP is used as a SG specific marker.

