## Supplementary figures and images for "The RNA demethylases ALKBH5 and FTO regulate translation of the ATF4 mRNA in sorafenib-treated hepatocarcinoma cells"

### Supplementary data 1

#### Supplementary data 2

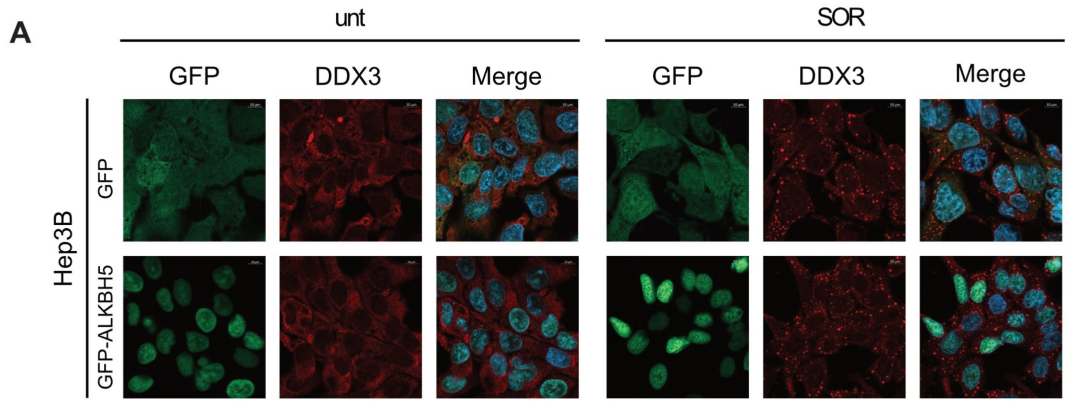


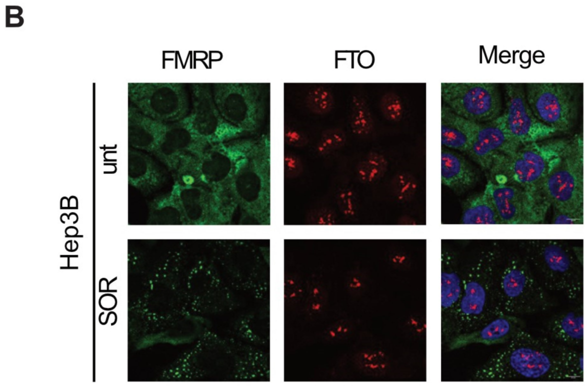

### Supplementary data 1

#### Supplementary data 1

A

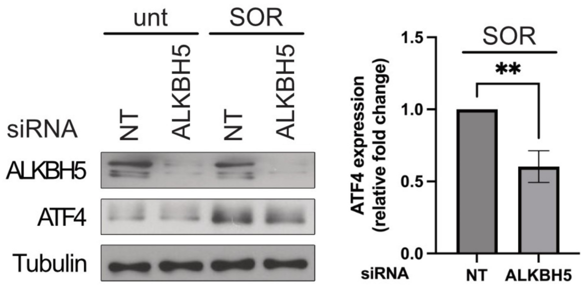


B

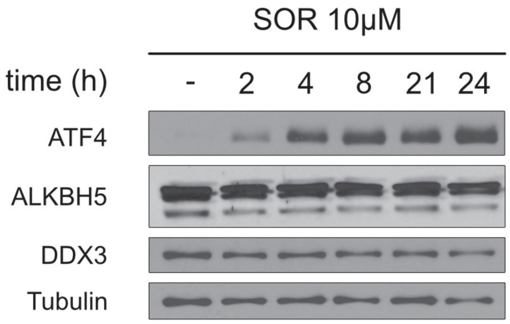
